# Genomic re-sequencing reveals mutational divergence across genetically engineered strains of model archaea

**DOI:** 10.1101/2024.08.08.607208

**Authors:** Andrew L. Soborowski, Rylee K. Hackley, Sungmin Hwang, Guangyin Zhou, Keely Dulmage, Peter Schönheit, Charles Daniels, Alexandre W. Bisson-Filho, Anita Marchfelder, Julie A. Maupin-Furlow, Thorsten Allers, Amy K. Schmid

## Abstract

Because archaea are the evolutionary ancestors of eukaryotes, archaeal molecular biology has been a topic of intense recent research. The hypersaline adapted archaeal species *Halobacterium salinarum* and *Haloferax volcanii* serve as important model organisms because facile tools enable genetic manipulation. As a result, the number of strains in circulation among the haloarchaeal research community has increased over the last few decades. However, the degree of genetic divergence and effects on genetic integrity during inter-lab transfers remain unclear. To address this question, we performed whole genome re-sequencing on a cross-section of wild-type, parental, and knockout strains in both model species. Integrating these data with existing repositories of re-sequencing data, we identify mutations that have arisen in a collection of 60 strains, sampled from 2 species across 8 different labs. Independent of sequencing, we construct strain lineages, identifying branch points and significant genetic effects in strain history. Combining this with our sequencing data, we identify small clusters of mutations that definitively separate lab strains. Additionally, an analysis of gene knockout strains suggests that roughly 1 in 3 strains currently in use harbors second-site mutations of potential phenotypic impact. Overall, we find that divergence among lab strains is thus far minimal, though as the archaeal research community continues to grow, careful strain provenance and genomic re-sequencing are required to keep inter-lab divergence to a minimum, prevent the compounding of mutations into fully independent lineages, and maintain the current high degree of reproducible research between lab groups in the haloarchaeal research community.

**Data Summary:** Novel sequencing data for this project was submitted to the National Center for Biotechnology Information (NCBI) Sequence Read Archive (SRA) and can be found under bioproject accession PRJNA1120443. SRA accessions for previously published sequencing data are available in supplementary table 1. R code for performing analysis and generating figures is available at https://github.com/andrew-soborowski/halophile_genome_resequencing.

**Impact Statement:** Archaea are important due to their shared evolutionary history with eukaryotes. As the archaeal research community grows, keeping track of the genetic integrity of archaeal strains of interest is necessary. In particular, routine genetic manipulations and the common practice of sharing strains between labs allow mutations to arise in lab stocks. If these mutations affect cellular processes, they may jeopardize the reproducibility of work between research groups and confound the results of future studies. In this work, we examine DNA sequences from 60 strains across two species of archaea. We identify shared and unique mutations occurring between and within strains. Independently, we trace the lineage of each strain, identifying which genetic manipulations lead to observed off-target mutations. While overall divergence across labs is minimal so far, our work highlights the need for labs to continue proper strain husbandry.

## Introduction

In recent decades, hypersaline-adapted archaea, hereafter referred to as haloarchaea, have emerged as leading model organisms in the study of archaeal genetics, genomics, and biochemistry [1–3]. Unique among archaeal lineages, this class has produced multiple genetically tractable organisms, facilitating both single-organism studies and class-wide comparisons. Haloarchaeal genomes, which tend to be organized similarly to bacteria, are characterized by genes co-regulated in operons and relatively short non-coding sequences. Considerable research has already been performed in two of these organisms, *Halobacterium salinarum* (*Hbt*.) and *Haloferax volcanii* (*Hfx*.), for which complete genome sequences [4, 5], counterselection-based gene knockout systems [6, 7], additional selectable markers [8], and overexpression plasmids [9], among other tools, have been developed [10]. Both species contain a single, circular chromosome containing the majority of their genes, in addition to a small number of additional replicons, pNRC100 and pNRC200 in *Hbt*. and pHV1, pHV2, pHV3, and pHV4 in *Hfx*. [4, 5].

These replicons, with the exception of the pHV2 plasmid, are referred to as mega-plasmids as they are large, contain essential genes, and contain functional origins of replications [11, 12]. However, the mega-plasmids are enriched for repetitive sequence and duplicated genes, complicating genomic assembly.

In particular, gene knockouts have become commonplace in the study of *Hbt*. and *Hfx*., owing to their relative simplicity given the power of existing toolkits. Current methodology in both species uses a selection / counterselection (“pop-in / pop-out”) approach [7]. The system exploits the uracil biosynthesis pathway by generating unmarked mutations in two steps of selection, similar to that used commonly in yeast genetics. This system facilitates a relatively easy process to generate gene knockouts using a uracil auxotroph strain as the parent from which subsequent mutant strains are derived. These strains are commonly used in both species, and we will refer to them as Δ*ura3* in *Hbt*. and Δ*pyrE* in *Hfx*. [6, 7].

At the conclusion of the pop-in pop-out procedure, it is crucial to verify the successful deletion of the gene because: (a) the procedure is equally likely to result in the parental and the desired knockout genotypes. If the desired knockout is deleterious, the parental strain revertant is even more likely; (b) secondary mutations arising during selection can alter the phenotype observed in any downstream experiments, confounding results; (c) haloarchaeal genomes are polyploid, harboring approximately 20 genome copies in a single cell during exponential phase [13, 14]. Polyploidy enables low-level wild-type gene copies to persist during and after selection if the gene is essential.

Genotypic verification is often performed using end-point PCR to amplify across the genomic region targeted for deletion, or a Southern blot to probe for the deleted sequence. However, recent research in our lab and others has shown that whole genome DNA re-sequencing (WGS) provides the sensitivity required to detect second-site mutations or low-level wild-type copies of the gene that remain in the polyploid genome below the level of detection by PCR [15–22]. Follow-up experiments to determine the phenotypic consequences of any detected second-site mutations are therefore important. For example, using WGS we have identified multiple second-site mutations, some with functional consequences (e.g. second-site mutations in the novel TF-coding gene *tbsP* suppresses lethality of *trmB* deletion under gluconeogenic conditions) [15–17]. In contrast, other second-site mutations did not affect the primary phenotype of interest [15]. Nonetheless, low levels of a wild-type copy of the gene of interest and/or second-site mutations undetectable with PCR or Southern blotting remain a primary concern in genetic manipulation of polyploid organisms. In a post-genomics world, WGS has become increasingly affordable, and thus increasingly suitable, for this task with the recent NovaSeq X promising a $200 human genome, about $2.00 per Gb of short reads. Long-read sequencing is also an option, trading cost and marginal accuracy for the ability to detect large-scale rearrangements and easily assemble across repetitive regions.

WGS is also an important method for periodic verification of the genetic integrity of the wild-type and parental strains. Although genetic analysis is typically done in relation to a published reference sequence, there is always a possibility of novel mutations arising in an individual lab’s stock due to either random drift or adaptation to the lab environment. As stock strains are continually passaged and transferred to new labs, the possibility increases for functional mutations to arise and propagate, confounding comparisons both to the static reference and comparisons made between labs. This concern is already well realized among labs studying common model organisms [23, 24] but this problem has yet to be addressed among haloarchaeal species. WGS via short-read next-generation sequencing therefore presents a relatively inexpensive, labor lite, and highly sensitive approach to address these concerns that are not detectable with PCR-based genetic verification of strains.

In this study, we report WGS data for 60 strains (51 of which have not previously been published) across both model haloarchaeal species with at least 29-fold coverage. We focused primarily on gene deletion strains without prior sequencing data to verify genetic backgrounds for use in further experiments, both in our lab and in the greater community. Additionally, we sequenced wild-type and uracil auxotroph parental strains across both species and sourced from many lab groups. This enabled the identification of mutations that have arisen at each step between the published reference strain and the final knockout, as well as any divergence that may have arisen between parental lab strains. With these data, we demonstrate that overall divergence between lab strains has thus far been minimal, though both lab-to-lab transfers and purposeful events generate mutations fixed between parental strains. Additionally, we identify cohorts of mutations fixed among each sampled strain in contrast to the published reference, as well as mutations fixed in distinct lineages. A survey across knockout strains in both species reveals a number of second-site mutations of potential phenotypic impact, highlighting the need for WGS verification when performing genetic manipulations.

## Methods

### Strain construction protocols

To construct knockout (KO) mutant strains of *Hfx. volcanii*, the double crossover selection method (“pop-in/pop-out”) was applied [7, 8, 25]. First, knockout (KO) plasmids were generated. Briefly, approximately 500 bp of the flanking regions of the sequence to be deleted were cloned into pTA131. For example, for Schmid lab vectors, pre-KO primers were used to generate PCR amplicons harboring the genomic regions flanking the target gene of interest (about 500 bp each, see primers for all labs and strains in S1 Table). These amplicons were then digested by appropriate restriction enzymes and inserted into the multiple-cloning sites of pTA131, resulting in “pre-KO” plasmids. For deletion strains generated in the Marchfelder laboratory (Δ*trh7*, Δ*tfb9*, Δ*tfb12*, Δ*arcR14* and Δ*arcR21*) the genomic region flanking each target gene (approx. 500 bp) were amplified as a single fragment by overlap PCR, resulting a single DNA fragment cloned into pTA131. These plasmids were utilized as templates to generate KO plasmids via reverse PCR with KO primers. For all labs, KO plasmids were transformed into *E. coli dam*-strain GM2163 prior to transformation of *Hfx. Volcanii* Δ*pyrE2* strain background (see S1 Table for the specific strain number used by each lab). As described in the Halohandbook (https://haloarchaea.com/halohandbook/), the polyethylene glycol (PEG) method was used for transformation followed by selection of pop-in transformants on Hv-CA plates without uracil. Pop-out transformants were subsequently selected on YPC plates supplemented with 50 *µ*g/mL 5-fluoroorotic acid (5-FOA). Clones were screened by PCR primers listed in S1 Table, then subjected to whole genome re-sequencing as detailed below. The Daniels lab *Hfx. volcanii pyrF* gene was deleted using the “pop-in/pop-out” strategy and checked using primers listed in S1 Table. For *Hbt. salinarum* KO strains, the same protocol was followed except that Δ*ura3* was the parent background [6], pNBK07 the cloning vector [26], and Complete Medium (CM) was used for routine growth (see [21] for recipe), and.

### Sources for strains reported in this study

For *Hbt*., strains were sourced from the Schmid and Baliga labs: a wildtype and 13 knockout strains from the Baliga lab [27–32], and a parental [6] with 6 knockout strains from the Schmid lab [16, 19, 21, 33, 34].

For *Hfx*., strains are sourced from seven labs: a wildtype from the Bisson lab, a parental strain from the Daniels lab, 6 knockout strains from the Marchfelder lab [35], a parental strain and 11 knockout strains from the Maupin-Furlow lab [7, 36], 2 knockout strains from the Schönheit lab [37, 38], a knockout and corresponding parental strain from the Peck lab [9, 39], and 14 knockout strains with the corresponding parent strain from the Schmid lab [7, 15, 17, 20, 40]. Note that parent strains for *Hfx*. originated from the Mevarech [7] and Allers labs [9], but were not resequenced here, see Results.

### Cultivation of strains, gDNA extraction, and sequencing

*Hfx. volcanii* strains (see S1 Table for complete strain list and additional sequencing details) were routinely grown by streaking from frozen glycerol stock onto solid YPC18% medium supplemented with 50 *µ*g uracil and grown at 42^*°*^C (see [9] for media formulation), except for *trmB* knockout strains, to which 1% glucose was added [28]. Genomic DNA (gDNA) was extracted from mid to late-expontential phase YPC18% liquid cultures using 25:24:1 phenol:chloroform:isoamyl alcohol and ethanol precipitation as described previously [19]. *Hbt*. strains were treated similarly except that the medium used was CM medium containing 50 *µ*g uracil (see [16] for media formulation). For both species, gDNA was subject to Illumina whole genome sequencing by the Duke Sequencing and Genomic Technologies Core Facility (see S1 Table for further information about sequencing platform).

### Identifying mutations

Before sequence alignment and variant calling, adapter sequence was trimmed and low-quality reads removed using trimgalore [41]. Sequences for each species were aligned to their respective reference genome (*Hbt*. at https://www.ncbi.nlm.nih.gov/datasets/genome/GCF_000006805.1/ accessed on December 19, 2022 and *Hfx*. at https://www.ncbi.nlm.nih.gov/datasets/genome/GCF_000025685.1/ accessed on September 12, 2022) using Bowtie2 [42] with mutations called by the microbial variant caller breseq using default parameters and run in clonal mode [43]. Mutational calls were tabulated using the supplemental tool GDtools [43]. Select variants of particular interest were confirmed by directly examining the read pileup using the Java script tool Integrative Genomics Viewer (IGV) [44].

### Mutation definition

For the purposes of this work, we define a mutation as a single, discrete event leading to an observable change in genome sequence compared to the reference. As both coverage depth and read length vary significantly across our samples (see S1 Table), we focus exclusively on mutations called with high confidence by breseq. For details on confidence thresholds and evidence types used to call mutations, please refer to the breseq manual: https://barricklab.org/twiki/pub/Lab/ToolsBacterialGenomeResequencing/documenta-tion/methods.html. We excluded: (a) low confidence calls, which can result from mixed non-clonal populations; (b) unassigned missing coverage and unassigned junction evidence, which can result from large-scale deletions or chromosomal rearrangements and/or highly repetitive sequences [43]; (c) certain high-quality calls with mutations in highly repetitive sequences. Mobile element mutations are identified solely as high-confidence mutations occurring inside IS elements and transposases (long-read sequencing data are required to resolve repeats and structural variation).

## Results

### Lineage tracing points to recent shared ancestry of most strains

To understand the genome-wide mutational landscape across laboratory domesticated haloarchaea, we performed short-read next-generation whole genome DNA re-sequencing (WGS) on a collection of 60 parental and gene knockout strains of *Hbt*. and *Hfx*., compiling a dataset of unique mutations as identified by the microbial mutation caller breseq [43] (see Methods). Since many of the strains in this collection were sourced from different labs, we first traced the lineage of each strain by combining a thorough literature review and personal correspondence. We traced commonly used parental strains back to the commercially available wild-type strain for both *Hbt*. (Fig 1A) and *Hfx*. (Fig 1B), identifying both genetic manipulations performed to create new parental strains and direct transfers of parental strains between labs. To distinguish between identical background strains sourced from different labs, here we label each parental strain with the lab’s initials. These labels correspond to the lab in which the strain was used in subsequent work, though we note that every strain but one was subsequently passaged and transferred to the Schmid lab for sequencing (S1 Table).

**Fig 1.**
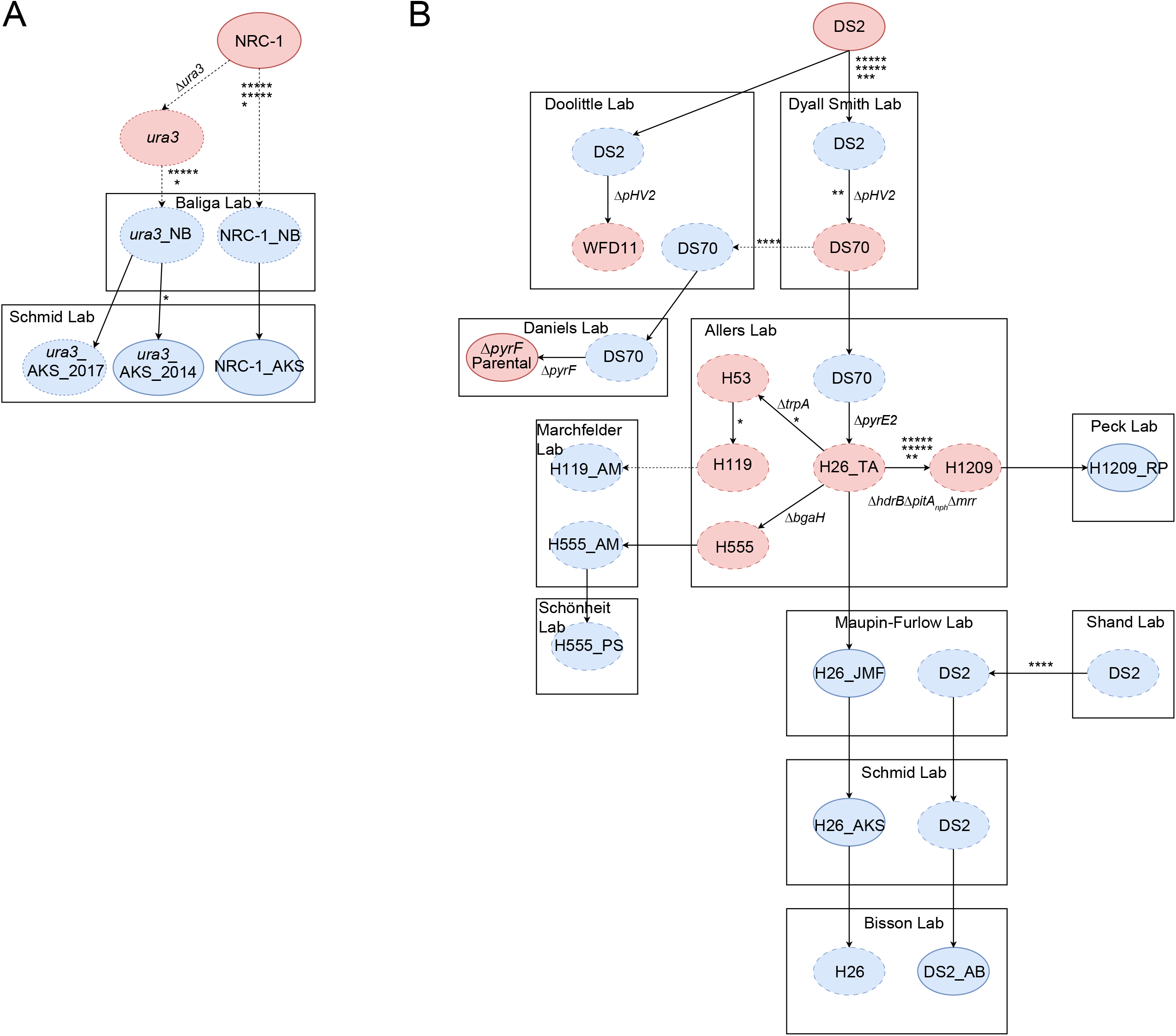
Lineage map of wildtype and parental strains. A. *Hbt*. and B. *Hfx*. diagrams of the lineage of sequenced and related parental and wild-type strains. Circles with solid outlines correspond to sequenced strains, whereas circles with dashed outlines correspond to known intermediate strains not sequenced here. Rectangles demarcate strains constructed within each lab. Full arrows correspond to known genetic manipulations (within rectangles) or transfers between labs (rectangle to rectangle). Dashed arrows refer to implied events with unknown intermediate steps. Sequenced parental or wild-type strains are assigned a unique postscript corresponding to their lab. Red circles correspond to the first instance of a given lineage, whereas blue circles depict strains transferred between labs. Asterisks enumerate the mutations relative to the closest sequenced ancestor. Unique identifiers for Δ*ura3*, Δ*pyrE2*, and Δ*pyrF* are given in S1 Table.

The *Hbt*. wild-type strain used in this study is NRC-1, first sequenced in 2000 and commercially available via ATCC (700922) [4]. Δ*ura3* is a uracil auxotroph and the sole *Hbt*. parental strain included in this study. Generated from NRC-1, Δ*ura3* was also developed in the year 2000 to enable the construction of unmarked deletion mutations [6]. Descendants of both strains were transferred to the Baliga lab, which we identify as NRC-1_NB and Δ*ura3*_NB. Δ*ura3*_NB was used as the parental strain to construct 13 knockout strains included in this study. Descendants of Δ*ura3*_NB were transferred to the Schmid lab in 2009 and later sequenced in 2014 (Δ*ura3*_AKS_2014) and 2017 (Δ*ura3*_AKS_2017). NRC-1_AKS, a descendant of the wild-type, was also sequenced in 2014. Δ*ura3*_AKS_2014 was used to construct 5 knockout strains included in this study and Δ*ura3*_AKS_2017 was used to construct 1 knockout strain included in this study.

We identified 11 mutations common across each *Hbt*. strain sequenced that differ from the published reference sequence [4], suggesting that these mutations arose: (a) before Δ*ura3* was generated from NRC-1; (b) in the specific NRC-1 isolate used to produce the reference genome (Fig 1A); or (c) from errors in the reference genome. Additionally, we identified 6 mutations common to every strain except NRC-1_AKS. As the most recent common ancestor of these strains is Δ*ura3*_NB and the mutations are undetected in the wild-type strain, it is most likely these arose alongside the knockout of *ura3* or during a previous transfer of the strain. An additional mutation was uniquely identified in all descendants of Δ*ura3*_AKS_2014, suggesting this particular mutation arose during transfer to the Schmid lab.

In the *Hfx*. lineage, the wild-type strain used in this study is DS2, sequenced in 2010 and available commercially via ATCC (29605) [5]. To enable genetic manipulation in *Hfx*., DS2 was cured of the pHV2 plasmid to give rise to the DS70 parental strain created in the Dyall Smith lab [45]. Subsequently, due to parallel efforts to improve DS70 for the construction of genetic knockout strains, the lineage of DS70 split, resulting in the Daniels lab Δ*pyrF* strain (Δ*pyrF* _CD) and the Allers lab Δ*pyrE* (H26_TA) strain. H26_TA serves as the ancestor of all knockout strains included in this study. Labs included in the current analysis either received their copy of H26 directly or secondarily (via an intermediate lab) from the Allers lab. Additionally, the Allers lab developed further descendants of H26_TA to facilitate genetic work, including H119, H555, and H1209 [8, 9]. We refer to H26, H119, H555, H1209, and Δ*pyrF* _CD as parental strains for the current analysis.

For *Hfx*., all knockout strains sequenced for this study are derived from five parental strains: the sequenced H26_AKS, H26_JMF, and H1209_RP, as well as the unsequenced H119_AM and H555_PS (Fig 1B, S1 Table). Each of these parental strains shares the Allers lab H26 strain as a common ancestor. In addition, wildtype DS2_AB and the Δ*pyrF* _CD parental strains were sequenced, although no knockouts derived from these are included here. DS2_AB is known to have been sequentially transferred between at least four labs, with the first known transfer occurring in 1999 from the Shand lab to the Maupin-Furlow lab [46].

Across these strains, we identified thirteen common mutations. Like in *Hbt*., these findings suggest that these mutations arose in the common ancestor of the DS2 lineage, in the specific DS2 strain used to construct the reference genome, or they represent errors in the assembly (Fig 1B). Additionally, we identified two mutations in every strain except DS2_AB, suggesting they arose after the last common ancestor but before the lineage split between H26_TA and Δ*pyrF* _CD. We identified four mutations unique to the Δ*pyrF* _CD lineage but none unique to the H26_TA lineage. Among the five parental strains from which the knockout strains were directly created, we identified no mutations in the H26_JMF, H26_AKS, or H555_PS lineages, though H26_AKS does carry two mutations not detected in any descendent strains, likely fixed only in the specific colony used for re-sequencing. The H119_AM lineage carries two mutations and the H1209_RP lineage carries twelve. Taken together, these results demonstrate that, for both species, mutations arise during the process of routine laboratory cultivation, generation of deletion strains, and inter-lab transfers.

### Class I mutations are shared across all strains tested

To further investigate mutations arising during inter-lab strain passages and within-lab genetic manipulation, we tabulated unique mutations (see Methods for how mutations were identified) appearing in each strain of the species *Hbt*. (Fig 2A) and *Hfx*. (Fig 2B). Across both species, mutations were separated into three classes. Class I: mutations detected in every strain of a given species, i.e. different from the published reference sequence. Class II: mutations unique to a parental strain and subsequently derived knockout strains. Class III: mutations unique to a specific knockout strain acquired during the process of strain construction. Additionally, mutations were further delineated between mobilome (mobile genetic elements, see Methods for definition) and non-mobilome related changes (Fig 3A,B). Detailed annotations for each reported mutation are given in S2 Table for *Hbt*. and S3 Table for *Hfx*.

**Fig 2.**
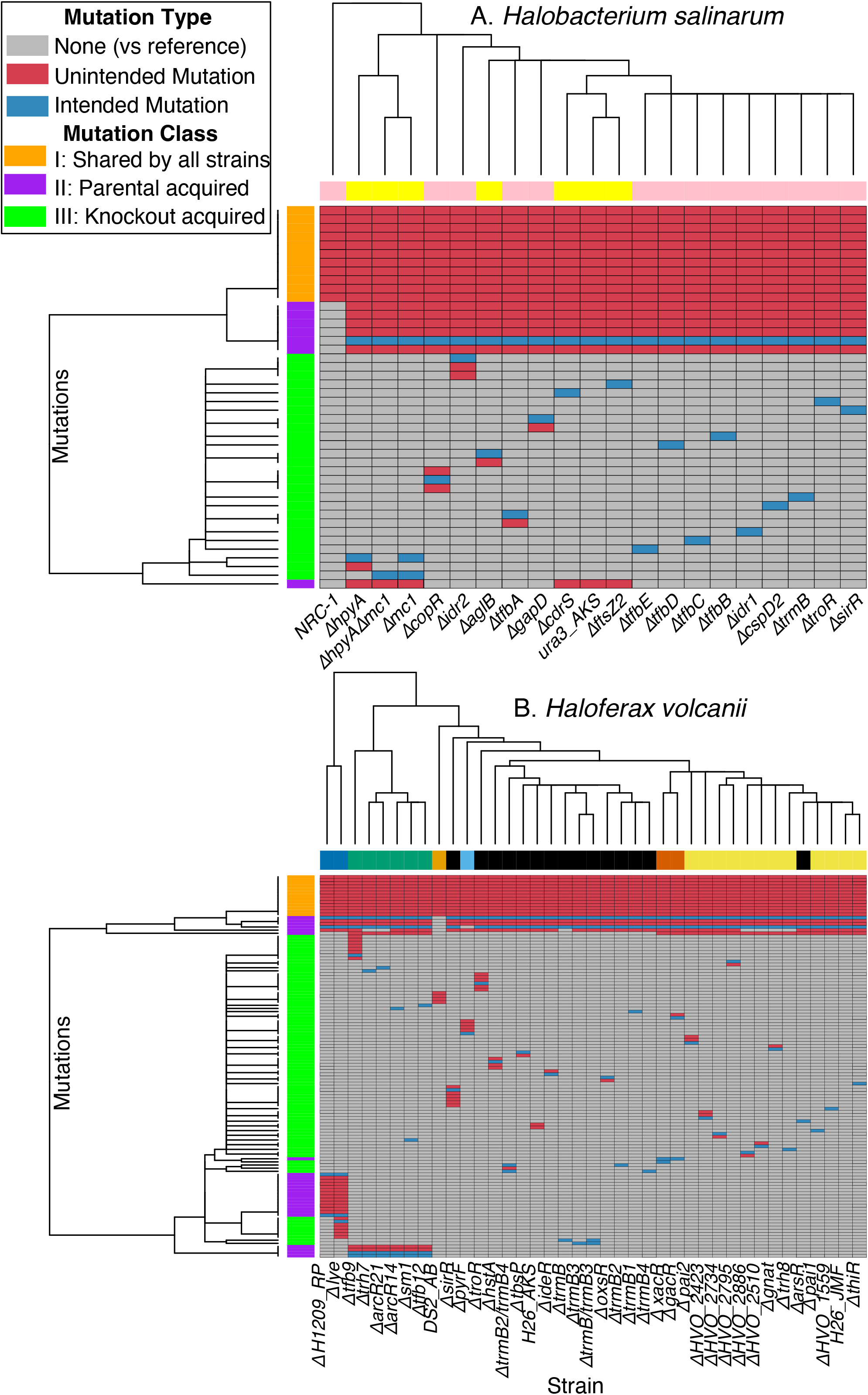
Heatmap of presence of mutation A. *Hbt*. and B. *Hfx*. Across both panels (A) and (B), gray entries correspond to mutations not present in the strain, blue entries correspond to experimenter-intended mutations that are present in the strain, and red entries correspond to mutations that are not intended to be made in the strain. Colors on the vertical bars represent the classes of mutations: orange represents Class I mutations shared by all strains (that differ from the published type strain [5]; purple, Class II, acquired in parental strain(s); green, Class III mutations, were acquired only in the knockout strains. Horizontal bars represent the lab of origin. For *Hbt*.: yellow, Schmid lab; pink, Baliga. For *Hfx*.: dark blue, Peck; green, Marchfelder; light orange, Bisson; black, Schmid; light blue, Daniels; dark orange, Schönheit, yellow, Maupin-Furlow. Clustering was performed on both strains and mutations with mutation clustering only on the presence or absence of the gene.

**Fig 3.**
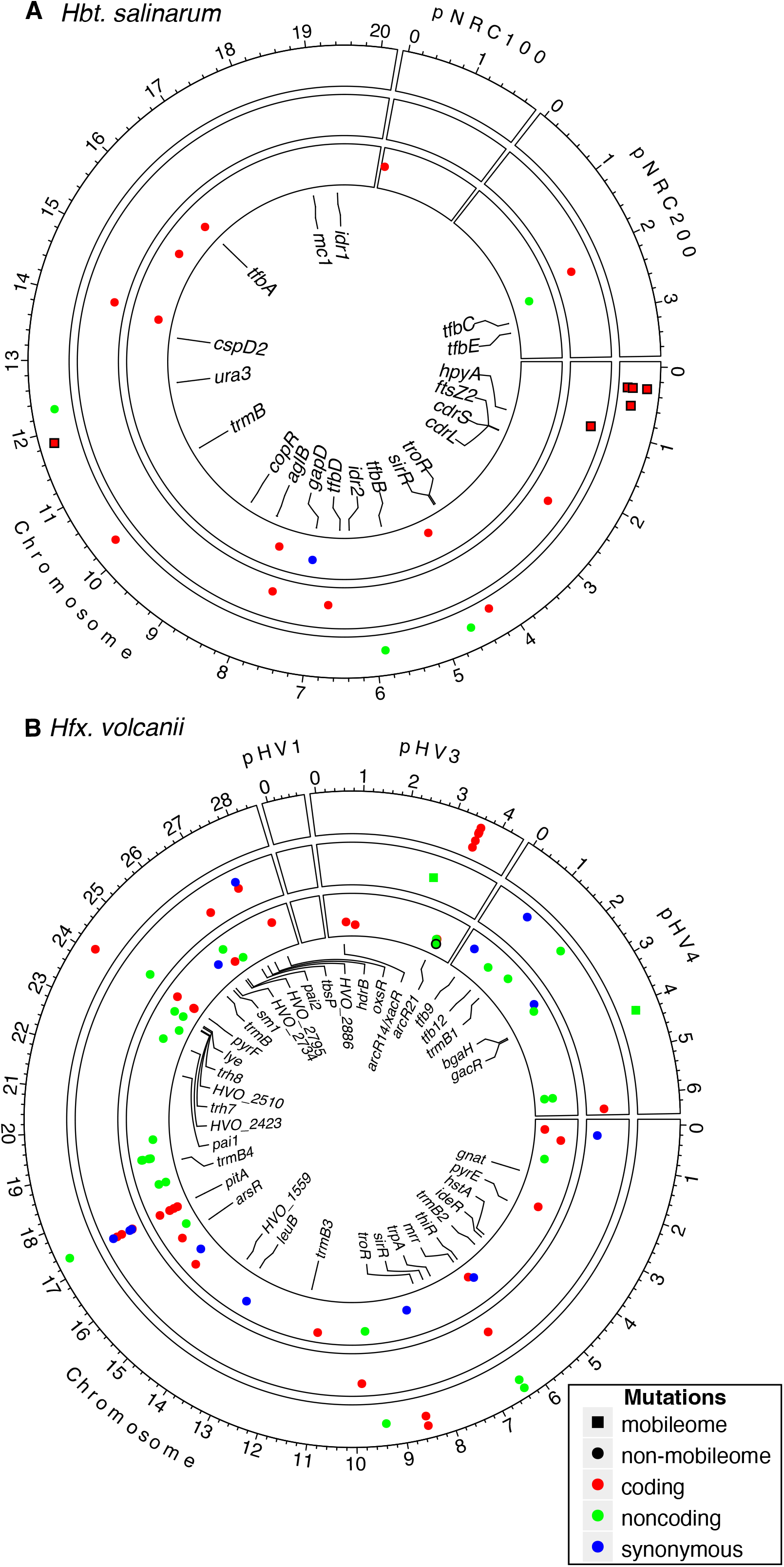
Genome plot of detected mutations. A. *Hbt*. and B. *Hfx*. Genomic distance is measured in Mbp. Each point corresponds to a unique mutation. Sequence features are: black, non-mobilome; red, coding region; green, non-coding regions. Mutation types: blue, synonymous mutations; black, red, and green, non-synonymous mutations; see also pictorial legend. Mutations in the mobilome are represented as squares. Points are localized to the chromosomal coordinate at the start of the corresponding mutation. Concentric rings correspond to mutation class, with the outer corresponding to class I, the middle class II, and the inner class III. Gene names correspond to intended knockouts.

For *Hbt*., 11 total mutations were detected in the first class of mutations (shared across strains, Fig 1A, Fig 2A, Fig 3A, [4]). Of these, six are located within mobile elements (Fig 3A). Four of these mobilome mutations cluster in the same gene, VNG_RS00125, predicted to encode an IS5 family transpose. The gene is located in a highly repetitive region of the genome, however, the mutations occur in sequence unique to the gene. Two mutations resulted in the deletion of VNG_0059H and VNG_1587H, annotated as IS4-like element ISH8B and IS5-like element ISH11, respectively, for which exist multiple identical and nearly identical duplicates in the genome. A closer examination revealed read troughs in the unique sequence directly upstream and downstream of each gene, consistent with a transposition event. Of the remaining five mutations, one may be of functional significance: a single nucleotide polymorphism (SNP) was detected in the coding region of VNG_1374G, which changes a serine to an arginine codon (S2 Table, Fig 3A). This gene is annotated as a PAS domain S-box protein which may be involved as a sensor module or transcriptional activator. A single nucleotide insertion was also detected in the intergenic region between VNG_1651H and VNG_1653H. The remaining three mutations correspond to the loss of a single base in a poly-C or poly-G tract region in a hypothetical protein and two pseudogenes. These are likely the result of sequencing errors (either here or in the originally published type strain genome sequence) given the technical challenges of sequencing homopolymer tracts [47]. Taken together, these results suggest the published NRC-1 reference sequence [4] still accurately represents strains currently in use. Moreover, many of the observed differences occur either in highly repetitive sequences or involve polynucleotide runs, which may be indicative of sequencing error or transposon movement with little impact on phenotype.

For *Hfx*., 13 mutations belong to the first class (shared across all strains, Fig 1B, Fig 2B). One predicted mutation, a SNP in HVO_2547 (changes L–>P in a 50S ribosomal protein) was previously characterized and determined to be an error in the reference genome [48]. Three mutations were detected as single base indels either in intergenic regions (between HVO_1027 and HVO_1028, and between HVO_3045 and HVO_1878), distant from transcription start sites, or within mobile elements (HVO_A0408, encoding an ISH3 family transposase) (Fig 3B, S3 Table)). These are unlikely to have functional consequences. The remaining 9 of these 13 mutations represented single base indels or SNPs that occur within or upstream of genes critical for metabolism and stress response: five immediately upstream or within the coding region of HVO_B0311 (XdhC/CoxL family protein), two 64 bp apart within HVO_0712 (shikimate dehydrogenase), and two 9 bp apart within the coding sequence of HVO_0938 (universal stress protein). Therefore, unlike in *Hbt*., the majority of these *Hfx*. class I mutations could have functional consequences in gene expression and/or protein function.

### Class II mutations were acquired in the parent strains during transfers or genetic manipulation

In *Hbt*., six class II mutations are shared in the strains descended from the Baliga lab Δ*ura3* strain but undetected in the ancestral NRC-1 strain (Fig 2A, [4]). One corresponds to the expected deletion of the *ura3* gene. One of the remaining five unexpected mutations involves the deletion of an ISH3-like element ISH27-2 family transposase (VNG_RS00465), which may have also deleted a small number of bases in a hypothetical protein 20 bases upstream (Fig 3A). Three mutations are SNPs inducing amino acid changes in three distinct genes: *trpB, gyrB*, and VNG_6283H, a hypothetical protein located on the pNRC200 plasmid. *trpB* codes for subunit beta of the tryptophan synthase protein complex, while *gyrB* is predicted to code for subunit beta of a DNA topoisomerase involved in DNA supercoiling [49, 50]. Finally, a seven bp deletion was detected in VNG_1896C (predicted ATP-binding protein) exclusively in strains Δ*ura3*_AKS_2014 and its descendants (Δ*hpyA*, Δ*mc1*, Δ*hpyA*Δ*mc1*, Δ*cdrS*, Δ*ftsZ2*). This suggests that the VNG_1869C mutation arose during the transfer of the strain from the Baliga lab to the Schmid lab. Taken together, these six mutations in the second class suggest the potential for phenotypic differences between derivative strains and their corresponding parent. However, to our knowledge, no phenotypic differences between these strains aside from the intended uracil auxotrophy of the Δ*ura3* strain have yet been observed or reported.

In *Hfx*., 30 total class II mutations were detected. One intergenic mutation was detected exclusively in reads over 150 bp and one pseudogene mutation was detected randomly in a highly repetitive region; both were excluded from further analysis. Four were shared between each strain derived from H26_TA (Fig 2B): (a) expected deletion of the pHV2 plasmid and *pyrE* (Fig 3B); and (b) one SNP in each of HVO_0032 and HVO_1080 (Fig 3B). In contrast, five mutations were specific to the Daniels lab Δ*pyrF* strain, necessarily arising either during transfer to the Daniels lab or during the knockout of the *pyrF* gene within the Daniels lab. These mutations include the expected deletion of *pyrF*, a SNP in *ldpA* (dihydrolipoyl dehydrogenase), and three mutations, including a frameshift in HVO_2576, a J-domain containing protein potentially involved in protein assembly and translocation. Additionally, this lineage also lacks pHV2 and carries the HVO_0032 and HVO_1080 mutations, suggesting they originated before the H26 and Δ*pyrF* lineages split.

Among the parental lineages descended from H26_TA (H1209_RP, H26_ JMF, H26_AKS, H555_PS, H119_AM), we detect 19 additional, lineage-specific mutations (Fig 1B, Fig 2B, Fig 3B). Five of these are expected, corresponding to the gene knockouts required to make the H119, H555, and H1209 strains. An additional four mutations were specific to the H1209_RP lineage, which may correspond to difficulty for breseq to properly recognize the replaced copy of *pitA* [9]. Of the remaining 10 mutations, three correspond to synonymous changes and one to an intergenic SNP (Fig 3B). Four nonsynonymous SNPs were detected in the H1209_RP lineage. Two nonsynonymous SNPs were detected in the H119_AM lineage, occurring in metallopeptidase HVO_A0634 and calcium/sodium antiporter HVO_0714. The mutation in HVO_A0634 was previously identified in H53, an intermediate between H119 and H26 [48]. No additional mutations were detected in the H26_ JMF, H26_AKS, or H555_PS lineages. Taken together, these mutations in both species highlight a growing, though as yet small, divergence between the parental strains at a genetic level with unknown phenotypic consequences as subsequent genetic manipulations are performed in their descendants.

### Class III mutations, acquired during construction of deletion strains, may lead to phenotypic consequences

In *Hbt*., a total of 19 knockout strains were sequenced, including 18 single knockouts and one double knockout. We identified a total of eight unique, unexpected mutations across these strains with no strain having more than two (Fig 2A). Each strain averages 0.42 mutations. Of these, six mutations alter a coding sequence, one is synonymous, and one occurs in an intergenic region (Fig 3A). Of the six coding mutations, three consist of mutations in hypothetical genes: a SNP in VNG_2183H, a 6 bp polynucleotide duplication in VNG_1026H, and an 87 bp deletion in VNG_0597H. The remaining coding mutations consist of an ATG-ATA start codon SNP in *nadA*, a SNP in transcriptional regulator *lrp*, and the complete elimination of the pNRC100 plasmid. Detailed descriptions of each mutation are given in S2 Table.

In *Hfx*., a total of 34 knockout strains were sequenced, 32 single knockouts and two double knockouts. We identified a total of 42 unique, unexpected mutations across these strains (Fig 2B). Each strain has at most seven mutations, though on average each strain has only 1.07. Of these 42 mutations, 12 alter coding sequences while 7 are synonymous and 23 occur in noncoding sequence (Fig 3B). Each coding mutation affects only a single gene and is either a SNP or a short (9-41 bp) deletion. Detailed descriptions of each mutation are given in S3 Table.

Overall, mutations in *Hbt*. knockout strains appeared in only 6 of the 19 (32%) strains sampled, although they are likely to lead to phenotypic consequences. Mutations appear more frequently in *Hfx*., appearing in 18 out of 34 (53%) strains. Mutation density is also higher in *Hfx*. at 1.07 mutations per strain versus only 0.42 mutations per strain in *Hbt*.. However, *Hfx*. tends to accumulate more noncoding and synonymous mutations, accumulating 0.35 coding mutations per strain compared to only 0.32 in *Hbt*. These coding mutations are the most likely to cause phenotypic impact and they affect a broad range of genes, including enzymes, transporters, transcription factors, and genes of unknown function.

## Discussion

This study provides a significant expansion in the available number of short-read whole genome DNA sequencing datasets available in the Haloarchaea, adding 51 novel strain sequences across *Hbt*. and *Hfx*. Analyzing these sequences alongside previously described sequences using the mutation caller breseq, we identify a population of mutations present in each of the species. Combining identified mutations with catalogued lineage information, we identify the time and intermediate strain in which each mutation likely arose. This allows the classification of mutations according to their prevalence across all sequenced strains, enabling both the analysis of extant mutations and their potential phenotypic impacts, as well as prediction of the type and frequency of mutations likely to arise in consort with common laboratory operations.

Global mutations occurring in every sequenced strain (class I) were detected 11 times in *Hbt*. and 13 times in *Hfx*. While it is difficult to conclude in which lineage these arose, three possible scenarios are possible: (a) mutations arose in the last common ancestor of every sequenced strain; (b) mutations arose independently in the single isolate used to generate the reference sequence; or (c) sequencing errors. In the case of both species, these mutations tend toward repetitive elements, noncoding changes, and multiple hits to the same gene. Parental mutations, occurring in distinct lineages among the sequenced strains, were detected in the uracil auxotroph lineages in both species, Δ*ura3* in *Hbt*. and H26 in *Hfx*., as well as many of the branching *Hfx*. parent strains derived from H26. Many of these changes tended to be coding, including every parental mutation in *Hbt*. Knockout mutations, occurring uniquely in single strains or derived double knockout strains, were detected in 32% of *Hbt*. strains and 51% of *Hfx*. genes. Both strains accumulated a similar number of coding mutations per knockout, though *Hfx*. strains were more likely to accumulate other synonymous or non-coding changes. This analysis highlights the importance of maintaining a permanent frozen stock within each lab to avoid accumulating mutations during routine experiments and inter-lab transfers.

In categorizing the mutations into three classes, we identify three experimental activities in which mutations commonly arise: parental strain generation, inter-lab transfers, and knockout strain generation. Background mutations are expected by chance. For example, the background mutation rate for *Hfx*. is estimated at 3.15 × 10^−10^ per site per generation [51]. However, these three lab experimental activities involve selection via population bottlenecks during colony selection steps. Bottlenecking is then compounded with selective pressure during strain generation, allowing both random and selected mutations to fix after the bottleneck step. As we only sequence a small number of strains compared to the large number of intermediates that may have existed, it is often difficult to conclude when a mutation became fixed. For example, in *Hbt*., a mutation present in Δ*ura3*_AKS_2014 and all its descendants but not Δ*ura3*_NB provides evidence that lab transfers can result in fixed mutations. However, the lack of mutational enrichment in strains transferred to the Schmid lab for sequencing, and thus undergoing an additional lab transfer step, vs. native Schmid lab strains that have not suggests that many of the mutations we observed likely arose during strain generation steps and not as a result of the transfer process. Regardless of mutational source, our analysis suggests that, while overall divergence between strains remains low, phenotypically impactful mutations can and do arise. This highlights the importance of WGS genotyping coupled with complementation experiments to ensure validity and reproducibility of the genotype-phenotype relationship in both intentionally generated mutants and primary lab stock.

Our use of short-read sequencing data in combination with breseq does impose two major limitations in this analysis. Breseq was chosen as the mutation caller in the study as it is the primary tool used in our lab and others for analyzing genomic DNA for mutations. Breseq has successfully called second-site mutations in recent genomic analyses across the tree of life [15–17, 52–54]. However, the first limitation imposed by our methodology is that the detection of large-scale rearrangements and mutations in highly repetitive regions is problematic, as it is difficult to place rearrangement junctions with short reads and uniquely map short reads in highly repeated sequences.

Previous work in *Hbt*. has already shown rearrangements among repeat sequence families to be fairly common: on the order of 10^*−*3^ potential arrangements occur per repetitive sequence family per generation [10]. Recent work in *Hfx*. has also demonstrated a high propensity for genomic rearangement, including division of the main chromosome into independent, self-sufficient replicons in the course of performing serial gene deletions [55]. While breseq attempts to predict genomic rearrangements and other large-scale events, predictions are of lesser quality and uncertain at boundaries. Future studies that incorporate long-read sequencing data, either exclusively or in tandem with short-read data, would be particularly fruitful in this regard, especially as the accuracy of these methods is rapidly improving [56].

The second limitation imposed by our methodology is a lack of identification of mutations that are transient in the population during transfer and strain construction. In theory, the population of cells should be clonal after each genetic bottleneck event, resulting in an entire population of genetically identical cells, such that sampling the DNA sequence of any sub-sample is equivalent to any other sub-sample. However, the cells are not static after the bottleneck event, where we expect some variation to arise, but not become fixed, in the final population. Additionally, both species are known to be highly polyploid, carrying up to 20 copies of the chromosome [57]. Heterozygous states transiently exist in the absence of selection but can endure in the presence of selection [25]. Problematically, from strictly looking at aggregated DNA sequences, it is not possible to differentiate intra-chromosomal differences versus intra-cellular differences, preventing the identification of heterozygous mutations and their possible phenotypic effects. Tracking such transient low-level variants over time is therefore an interesting avenue to pursue in future research.

Supported by independent lineage tracing, we demonstrate that functional divergence between lab strains has thus far remained minimal, though mutations of potential phenotypic impact have begun to arise as strains are continually transferred and manipulated by different lab groups. Additionally, we demonstrate a consistent accumulation of coding mutations in knockout strains in both species. Taken together, our results highlight the need for vigilance when working with haloarchaeal strains.

Recommendations include minimizing events that may lead to strain divergence, obtaining their strains from a common, ancestral source, and verifying strain integrity via whole-genome re-sequencing and/or complementation assays whenever feasible when confirming the genotype and phenotype of a novel deletion strain.

## Supporting information

**S1 Table. Strain information table**.

**S2 Table. *Hbt*. mutation table**.

**S3 Table. *Hfx*. mutation table**.

## Funding Information

AB is a Pew Scholar in the Biomedical Sciences, supported by The Pew Charitable Trusts. This work was also supported by grants from the National Science Foundation MCB 1936024 and 1651117 to AKS. The funders had no role in study design, data collection and analysis, decision to publish, or preparation of the manuscript.

## Acknowledgments

The authors thank Sierra Watson, Mar Martinez-Pastor, Cindy Darnell, and Angie Vreugdenil-Hayslette for preparing gDNA for sequencing many of the included strains. The Marchfelder lab thanks the students of our Master practical course 2014 and our Bachelor and Erasmus students Nadja Raab, Jasmin Niederhofer, Stefan Werner and Tereza Vavrdova for help with generating the deletion strains Δ*trh7*, Δ*tfb9*, Δ*tfb12*, Δ*arcR14* and Δ*arcR21*. We thank the Duke Sequencing and Genomic Technologies core facility for their technical support with sequencing. Thank you to the Baliga lab for the transfer of *Hbt*. strains to the Schmid lab in 2009. We also thank the Schmid lab members for their support and critical feedback on the manuscript. AB is a Pew Scholar in the Biomedical Sciences, supported by The Pew Charitable Trusts. This work was supported by grants from the National Science Foundation MCB 1936024 and 1651117 to AKS.

## Conflicts of Interest

The authors declare that there are no conflicts of interest.

## Notes

https://github.com/andrew-soborowski/halophile_genome_resequencing

### Competing Interest Statement

The authors have declared no competing interest.

## References

1. Leigh JA, Albers SV, Atomi H, Allers T. Model organisms for genetics in the domain Archaea: methanogens, halophiles, Thermococcales and Sulfolobales. FEMS Microbiology Reviews. 2011;35(4):577–608. doi:10.1111/j.1574-6976.2011.00265.x.

2. Soppa J. From genomes to function: haloarchaea as model organisms. Microbiology. 2006;152(3):585–590. doi:10.1099/mic.0.28504-0.

3. Pérez-Arnaiz P, Dattani A, Smith V, Allers T. Haloferax volcanii—a model archaeon for studying DNA replication and repair. Open Biology. 2020;10(12):200293. doi:10.1098/rsob.200293.

4. Ng WV, Kennedy SP, Mahairas GG, Berquist B, Pan M, Shukla HD, et al. Genome sequence of Halobacterium species NRC-1. Proceedings of the National Academy of Sciences. 2000;97(22):12176–12181. doi:10.1073/pnas.190337797.

5. Hartman AL, Norais C, Badger JH, Delmas S, Haldenby S, Madupu R, et al. The Complete Genome Sequence of Haloferax volcanii DS2, a Model Archaeon. PLoS ONE. 2010;5(3):e9605. doi:10.1371/journal.pone.0009605.

6. Peck RF, DasSarma S, Krebs MP. Homologous gene knockout in the archaeon Halobacterium salinarum with ura3 as a counterselectable marker. Molecular Microbiology. 2000;35(3):667–676. doi:10.1046/j.1365-2958.2000.01739.x.

7. Bitan-Banin G, Ortenberg R, Mevarech M. Development of a Gene Knockout System for the Halophilic Archaeon Haloferax volcanii by Use of the pyrE Gene. Journal of Bacteriology. 2003;185(3):772–778. doi:10.1128/JB.185.3.772-778.2003.

8. Allers T, Ngo HP, Mevarech M, Lloyd RG. Development of additional selectable markers for the halophilic archaeon Haloferax volcanii based on the leuB and trpA genes. Applied and Environmental Microbiology. 2004;70(2):943–953. doi:10.1128/AEM.70.2.943-953.2004.

9. Allers T, Barak S, Liddell S, Wardell K, Mevarech M. Improved Strains and Plasmid Vectors for Conditional Overexpression of His-Tagged Proteins in Haloferax volcanii. Applied and Environmental Microbiology. 2010;76(6):1759–1769. doi:10.1128/AEM.02670-09.

10. Sapienza C, Rose MR, Doolittle WF. High-frequency genomic rearrangements involving archaebacterial repeat sequence elements. Nature. 1982;299(5879):182–185. doi:10.1038/299182a0.

11. Coker JA, DasSarma P, Capes M, Wallace T, McGarrity K, Gessler R, et al. Multiple Replication Origins of Halobacterium sp. Strain NRC-1: Properties of the Conserved orc7-Dependent oriC1. Journal of Bacteriology. 2009;191(16):5253–5261. doi:10.1128/JB.00210-09.

12. Norais C, Hawkins M, Hartman AL, Eisen JA, Myllykallio H, Allers T. Genetic and Physical Mapping of DNA Replication Origins in Haloferax volcanii. PLOS Genetics. 2007;3(5):e77. doi:10.1371/journal.pgen.0030077.

13. Breuert S, Allers T, Spohn G, Soppa J. Regulated Polyploidy in Halophilic Archaea. PLOS ONE. 2006;1(1):e92. doi:10.1371/journal.pone.0000092.

14. Ludt K, Soppa J. Polyploidy in halophilic archaea: regulation, evolutionary advantages, and gene conversion. Biochemical Society Transactions. 2019;47(3):933–944. doi:10.1042/BST20190256.

15. Martinez Pastor M, Sakrikar S, Hwang S, Hackley R, Soborowski A, Maupin-Furlow J, et al. TroR is the primary regulator of the iron homeostasis transcription network in the halophilic archaeon Haloferax volcanii. Nucleic Acids Research. 2024;52(1):125–140. doi:10.1093/nar/gkad997.

16. Darnell CL, Zheng J, Wilson S, Bertoli RM, Bisson-Filho AW, Garner EC, et al. The Ribbon-Helix-Helix Domain Protein CdrS Regulates the Tubulin Homolog ftsZ2 To Control Cell Division in Archaea. mBio. 2020;11(4):10.1128/mbio.01007–20. doi:10.1128/mbio.01007-20.

17. Hackley RK, Hwang S, Herb JT, Bhanap P, Lam K, Vreugdenhil A, et al. TbsP and TrmB jointly regulate gapII to influence cell development phenotypes in the archaeon Haloferax volcanii. Molecular Microbiology. 2024;n/a(n/a). doi:10.1111/mmi.15225.

18. Hackley RK, Vreugdenhil-Hayslette A, Darnell CL, Schmid AK. A conserved transcription factor controls gluconeogenesis via distinct targets in hypersaline-adapted archaea with diverse metabolic capabilities. PLOS Genetics. 2024;20(1):e1011115. doi:10.1371/journal.pgen.1011115.

19. Zaretsky M, Darnell CL, Schmid AK, Eichler J. N-Glycosylation Is Important for Halobacterium salinarum Archaellin Expression, Archaellum Assembly and Cell Motility. Frontiers in Microbiology. 2019;10.

20. Sakrikar S, Hackley RK, Martinez-Pastor M, Darnell CL, Vreugdenhil A, Schmid AK. The Hypersaline Archaeal Histones HpyA and HstA Are DNA Binding Proteins That Defy Categorization According to Commonly Used Functional Criteria. mBio. 2023;14(2):e03449–22. doi:10.1128/mbio.03449-22.

21. Sakrikar S, Schmid A. An archaeal histone-like protein regulates gene expression in response to salt stress. Nucleic Acids Research. 2021;49(22):12732–12743. doi:10.1093/nar/gkab1175.

22. Schiller H, Hong Y, Kouassi J, Rados T, Kwak J, DiLucido A, et al. Identification of structural and regulatory cell-shape determinants in Haloferax volcanii. Nature Communications. 2024;15(1):1414. doi:10.1038/s41467-024-45196-0.

23. Liu B, Eydallin G, Maharjan RP, Feng L, Wang L, Ferenci T. Natural Escherichia coli isolates rapidly acquire genetic changes upon laboratory domestication. Microbiology (Reading, England). 2017;163(1):22–30. doi:10.1099/mic.0.000405.

24. Zeigler DR, Prágai Z, Rodriguez S, Chevreux B, Muffler A, Albert T, et al. The origins of 168, W23, and other Bacillus subtilis legacy strains. Journal of Bacteriology. 2008;190(21):6983–6995. doi:10.1128/JB.00722-08.

25. Dattani A, Sharon I, Shtifman-Segal E, Robinzon S, Gophna U, Allers T, et al. Differences in homologous recombination and maintenance of heteropolyploidy between Haloferax volcaniiand Haloferax mediterranei. G3 Genes|Genomes|Genetics. 2023;13(4):jkac306. doi:10.1093/g3journal/jkac306.

26. Wilbanks EG, Larsen DJ, Neches RY, Yao AI, Wu CY, Kjolby RAS, et al. A workflow for genome-wide mapping of archaeal transcription factors with ChIP-seq. Nucleic Acids Research. 2012;40(10):e74. doi:10.1093/nar/gks063.

27. Schmid AK, Pan M, Sharma K, Baliga NS. Two transcription factors are necessary for iron homeostasis in a salt-dwelling archaeon. Nucleic Acids Research. 2011;39(7):2519–2533. doi:10.1093/nar/gkq1211.

28. Schmid AK, Reiss DJ, Pan M, Koide T, Baliga NS. A single transcription factor regulates evolutionarily diverse but functionally linked metabolic pathways in response to nutrient availability. Molecular Systems Biology. 2009;5(1):282. doi:10.1038/msb.2009.40.

29. Kaur A, Pan M, Meislin M, Facciotti MT, El-Gewely R, Baliga NS. A systems view of haloarchaeal strategies to withstand stress from transition metals. Genome Research. 2006;16(7):841–854. doi:10.1101/gr.5189606.

30. Martinez-Pastor M, Lancaster WA, Tonner PD, Adams M, Schmid AK. A transcription network of interlocking positive feedback loops maintains intracellular iron balance in archaea. Nucleic Acids Research. 2017;45(17):9990–10001. doi:10.1093/nar/gkx662.

31. Facciotti MT, Reiss DJ, Pan M, Kaur A, Vuthoori M, Bonneau R, et al. General transcription factor specified global gene regulation in archaea. Proceedings of the National Academy of Sciences. 2007;104(11):4630–4635. doi:10.1073/pnas.0611663104.

32. Bonneau R, Facciotti MT, Reiss DJ, Schmid AK, Pan M, Kaur A, et al. A Predictive Model for Transcriptional Control of Physiology in a Free Living Cell. Cell. 2007;131(7):1354–1365. doi:10.1016/j.cell.2007.10.053.

33. Dulmage KA, Todor H, Schmid AK. Growth-Phase-Specific Modulation of Cell Morphology and Gene Expression by an Archaeal Histone Protein. mBio. 2015;6(5):10.1128/mbio.00649–15. doi:10.1128/mbio.00649-15.

34. Darnell CL, Tonner PD, Gulli JG, Schmidler SC, Schmid AK. Systematic Discovery of Archaeal Transcription Factor Functions in Regulatory Networks through Quantitative Phenotyping Analysis. mSystems. 2017;2(5):10.1128/msystems.00032–17. doi:10.1128/msystems.00032-17.

35. Maier LK, Benz J, Fischer S, Alstetter M, Jaschinski K, Hilker R, et al. Deletion of the Sm1 encoding motif in the lsm gene results in distinct changes in the transcriptome and enhanced swarming activity of Haloferax cells. Biochimie. 2015;117:129–137. doi:10.1016/j.biochi.2015.02.023.

36. Hwang S, Cordova B, Abdo M, Pfeiffer F, Maupin-Furlow JA. ThiN as a Versatile Domain of Transcriptional Repressors and Catalytic Enzymes of Thiamine Biosynthesis. Journal of Bacteriology. 2017;199(7):10.1128/jb.00810–16. doi:10.1128/jb.00810-16.

37. Johnsen U, Sutter JM, Schulz AC, Tästensen JB, Schönheit P. XacR – a novel transcriptional regulator of D-xylose and L-arabinose catabolism in the haloarchaeon Haloferax volcanii. Environmental Microbiology. 2015;17(5):1663–1676. doi:10.1111/1462-2920.12603.

38. Tästensen JB, Johnsen U, Reinhardt A, Ortjohann M, Schönheit P. D-galactose catabolism in archaea: operation of the DeLey–Doudoroff pathway in Haloferax volcanii. FEMS Microbiology Letters. 2020;367(1):fnaa029. doi:10.1093/femsle/fnaa029.

39. Peck RF, Pleşa AM, Graham SM, Angelini DR, Shaw EL. Opsin-Mediated Inhibition of Bacterioruberin Synthesis in Halophilic Archaea. Journal of Bacteriology. 2017;199(21):10.1128/jb.00303–17. doi:10.1128/jb.00303-17.

40. Mondragon P, Hwang S, Kasirajan L, Oyetoro R, Nasthas A, Winters E, et al. TrmB Family Transcription Factor as a Thiol-Based Regulator of Oxidative Stress Response. mBio. 2022;13(4):e00633–22. doi:10.1128/mbio.00633-22.

41. Krueger F. Trim Galore; 2023. Available from: https://github.com/FelixKrueger/TrimGalore.

42. Langmead B, Salzberg SL. Fast gapped-read alignment with Bowtie 2. Nature Methods. 2012;9(4):357–359. doi:10.1038/nmeth.1923.

43. Deatherage DE, Barrick JE. Identification of mutations in laboratory evolved microbes from next-generation sequencing data using breseq. Methods in molecular biology (Clifton, NJ). 2014;1151:165–188. doi:10.1007/978-1-4939-0554-6_12.

44. Robinson JT, Thorvaldsdóttir H, Winckler W, Guttman M, Lander ES, Getz G, et al. Integrative Genomics Viewer. Nature biotechnology. 2011;29(1):24–26. doi:10.1038/nbt.1754.

45. Wendoloski D, Ferrer C, Dyall-Smith ML. A new simvastatin (mevinolin)-resistance marker from Haloarcula hispanica and a new Haloferax volcanii strain cured of plasmid pHV2. Microbiology. 2001;147(4):959–964. doi:10.1099/00221287-147-4-959. The GenBank accession number for the sequence reported in this paper is AF123438.

46. Wilson HL, Aldrich HC, Maupin-Furlow J. Halophilic 20S Proteasomes of the Archaeon Haloferax volcanii: Purification, Characterization, and Gene Sequence Analysis. Journal of Bacteriology. 1999;181(18):5814–5824.

47. Stoler N, Nekrutenko A. Sequencing error profiles of Illumina sequencing instruments. NAR Genomics and Bioinformatics. 2021;3(1):lqab019. doi:10.1093/nargab/lqab019.

48. Hawkins M, Malla S, Blythe MJ, Nieduszynski CA, Allers T. Accelerated growth in the absence of DNA replication origins. Nature. 2013;503(7477):544–547. doi:10.1038/nature12650.

49. Coker JA, DasSarma P, Kumar J, Müller JA, DasSarma S. Transcriptional profiling of the model Archaeon Halobacterium sp. NRC-1: responses to changes in salinity and temperature. Saline Systems. 2007;3:6. doi:10.1186/1746-1448-3-6.

50. Duprey A, Groisman EA. The regulation of DNA supercoiling across evolution. Protein Science : A Publication of the Protein Society. 2021;30(10):2042–2056. doi:10.1002/pro.4171.

51. Kucukyildirim S, Behringer M, Williams EM, Doak TG, Lynch M. Estimation of the Genome-Wide Mutation Rate and Spectrum in the Archaeal Species Haloferax volcanii. Genetics. 2020;215(4):1107–1116. doi:10.1534/genetics.120.303299.

52. Liu G, Catacutan DB, Rathod K, Swanson K, Jin W, Mohammed JC, et al. Deep learning-guided discovery of an antibiotic targeting Acinetobacter baumannii. Nature Chemical Biology. 2023;19(11):1342–1350. doi:10.1038/s41589-023-01349-8.

53. Stabryla LM, Johnston KA, Diemler NA, Cooper VS, Millstone JE, Haig SJ, et al. Role of bacterial motility in differential resistance mechanisms of silver nanoparticles and silver ions. Nature Nanotechnology. 2021;16(9):996–1003. doi:10.1038/s41565-021-00929-w.

54. Federici S, Kredo-Russo S, Valdés-Mas R, Kviatcovsky D, Weinstock E, Matiuhin Y, et al. Targeted suppression of human IBD-associated gut microbiota commensals by phage consortia for treatment of intestinal inflammation. Cell. 2022;185(16):2879–2898.e24. doi:10.1016/j.cell.2022.07.003.

55. Ausiannikava D, Mitchell L, Marriott H, Smith V, Hawkins M, Makarova KS, et al. Evolution of Genome Architecture in Archaea: Spontaneous Generation of a New Chromosome in Haloferax volcanii. Molecular Biology and Evolution. 2018;35(8):1855–1868. doi:10.1093/molbev/msy075.

56. Manuel JG, Heins HB, Crocker S, Neidich JA, Sadzewicz L, Tallon L, et al. High Coverage Highly Accurate Long-Read Sequencing of a Mouse Neuronal Cell Line Using the PacBio Revio Sequencer; 2023. Available from: 10.1101/2023.06.06.543940v1.

57. Zerulla K, Soppa J. Polyploidy in haloarchaea: advantages for growth and survival. Frontiers in Microbiology. 2014;5. doi:10.3389/fmicb.2014.00274.

